# CoVar: A generalizable machine learning approach to identify the coordinated regulators driving variational gene expression

**DOI:** 10.1101/2023.01.12.523808

**Authors:** Satyaki Roy, Shehzad Z. Sheikh, Terrence S. Furey

**Affiliations:** Department of Genetics, University of North Carolina, Chapel Hill, USA; Departments of Medicine and Genetics, Center for Gastrointestinal Biology and Disease, University of North Carolina, Chapel Hill, USA; Departments of Genetics and Biology, Center for Gastrointestinal Biology and Disease, University of North Carolina, Chapel Hill, USA

**Keywords:** Disease biology, Machine learning, Network inference, Gene expression

## Abstract

Network inference is used to model transcriptional, signaling, and metabolic interactions among genes, proteins, and metabolites that identify biological pathways influencing disease pathogenesis. Advances in machine learning (ML)-based inference models exhibit the predictive capabilities of capturing latent patterns in genomic data. Such models are emerging as an alternative to the statistical models identifying causative factors driving complex diseases. We present CoVar, an inference framework that builds upon the properties of existing inference models, to find the central genes driving perturbed gene expression across biological states. We leverage ML-based network inference to find networks that capture the strength of regulatory interactions. Our model first pinpoints a subset of genes, termed variational, whose expression variabilities typify the differences in network connectivity between the control and perturbed data. Variational genes, by being differentially expressed themselves or possessing differentially expressed neighbor genes, capture gene expression variability. CoVar then creates subnetworks comprising variational genes and their strongly connected neighbor genes and identifies core genes central to these subnetworks that influence the bulk of the variational activity. Through the analysis of yeast expression data perturbed by the deletion of the mitochondrial genome, we show that CoVar identifies key genes not found through independent differential expression analysis.

## 1. Introduction

The advent of high-throughput genomic data acquisition techniques has generated immense interest in the statistical and data-driven analysis of biological and biochemical interactions [1]. One such approach, network inference, attempts to identify network topologies that capture the “interactome”, defined as the set of direct or indirect molecular interactions, of the biological system [2, 3]. It employs a range of computational models, such as Bayesian, autoregression, and differential equations, to answer a wide range of questions ranging from basic cell biology to disease pathogenesis. The efficacy of the resultant biological networks largely depends on the ability of the underlying computational models to capture the complex dynamics among the entities determined by nonlinear and stochastic interactions [4].

Efforts to carry out network analysis of gene expression data show that the genes acting as key players in biological processes are often highly connected nodes in the corresponding biological networks [5, 6]. Thus, a measure of the functional significance of a gene is its ability to have paths that reach many genes in the network. On the other hand, there have been attempts to group genes into modules based on similarity in expression profiles and infer the aggregate biological function of these modules [7, 8]. The computational methods to identify such modules, such as the Weighted Correlation Network Analysis [9] and Sub-Network Enrichment Analysis algorithm (SNEA), reveal subnetworks with potential regulators as well as target regulated genes, often using information from known databases of genetic interactions, where multiple regulator genes can coordinate to influence several cellular processes [10]. Thus, coordination and reachability are two necessary facets of biological network analysis.

There are predominantly two classes of network inference approaches. The first is to learn the network structure completely from the genomic data. For instance, co-expression networks are constructed by connecting genes with highly similar or correlated gene expression profiles [11]. Liu et al. attempted to find accurate protein-protein interaction networks, by adding and removing existing links based on topological and semantic similarity between proteins [12]. Werhli et al. presented a graphical Gaussian model that builds a conditional independence relationship among genes under the assumption that the data follow multivariate Gaussian distribution [13] and Murphy et al. proposed a directed acyclic graph-based where directed links capture the interaction among genes [14]. Steuer et al. used mutual information on the expression profile of genes as a measure of the regulatory interaction between them [15]. The second class of network inference leverage known associations to make inferences. Christley et al. incorporated prior transcription factor-gene interaction into gene network inference [16]. Siahpirani et al. introduced a regulatory network inference algorithm that integrates expression information from auxiliary datasets [17]. Li et al. included prior biological knowledge in gene networks by adding associations between gene pairs based on gene ontology, protein-protein interaction, and gene regulatory networks [18].

Key genes responsible for altered cellular activity between biological states are often predicted by determining individual genes whose expression profiles are significantly altered, but these changes may be better analyzed at a network level to capture effects across multiple genes and pathways. To identify differences between networks from two different datasets or conditions, Zhou et al. presented an approach that creates a joint inference of two gene regulatory networks (GRNs) by fusion of gene expression and genetic perturbation data to facilitate the identification of the differential GRN under two conditions [19]. Tu et al. conceived a differential network where differential edges exist only if at least one of the two involved genes is differentially expressed. They showed that the hub nodes estimated by the model have significant biological roles [20]. Nitsch et al. introduced a network analysis tool that identifies disease-causing genes under the assumption that such genes are surrounded by differentially expressed genes [21]. Network centrality measures, such as degree, betweenness and page rank centrality, are often used to identify the most well-connected genes, which, when removed from the network, can disrupt key functional properties of the biological processes being modeled [22, 23]. Langfelder et al. utilized network clustering analysis to identify modules of interrelated genes with similar functional roles [9], while Kogelman et al. proposed a probabilistic graphical model to elucidate transcriptomic pathways and regulators genes involved in disease pathogenesis [24]. Khwaja et al. and He et al. attempted to eliminate the strong and dominant clusters identified by existing community detection algorithms to be able to identify weak or hidden clusters within networks [25, 26].

In this work, we present a unified approach, called CoVar, that builds upon some of the key features of existing network inference models to identify the central genes driving perturbation in gene expression datasets. An important difference is that existing inference methods rely on modules of correlated gene expression profiles or on prior associations to guide identification of central hub genes. In contrast, CoVar leverages a set of genes, termed *variational*, that exhibit a significant relative difference in interactions with neighbor genes across the control and perturbed expression datasets. Thus, CoVar focuses on *changes in expression relationships*, rather than commonalities in expression profiles, to infer modules of coordinated regulators driving perturbation. CoVar first creates directed networks independently in each condition using only the gene expression data. It employs a machine learning-based network inference approach called GENIE3 [27, 28] that captures the strength of direct or indirect interactions between genes and helps identify the variational genes. The variational genes may themselves be differentially expressed or may possess neighbor genes that are differentially expressed, thereby capturing the relative variability in gene expression due to perturbation. Based on this principle, CoVar creates a *nearest neighbor network* that includes the variational genes and their strongly connected neighbor genes. Within this network, distinct subnetworks or modules that display a high degree of intra-module connectivity and a low or no degree of inter-module connectivity are then determined. Finally, a subset of genes, termed core genes, are identified within each module that exhibits both coordination through strong mutual interaction and reachability with short paths to nearest neighbor network genes within the module. These modules suggest key biological processes being altered in the perturbed state with their core genes playing an essential role in facilitating their change in activity. We demonstrate the features and the utility of CoVar using both simulated data as well as data from yeast experiments focusing on the effects of mitochondrial genome depletion.

## Methods

### 2.1 RNA-seq data preprocessing

CoVar takes as input results from two sets of genome-wide expression experiments, where one set of experiments represent a perturbation in relation to the second set of control experiments. Data from each experiment is represented as *S* × *N* expression matrix ***X*** with *S* samples and *N* genes, where rows are samples and columns are gene names, i.e., ***X**_i,u_* the expression value of gene *u* in sample *i* (see Figure 1).

**Figure 1:**
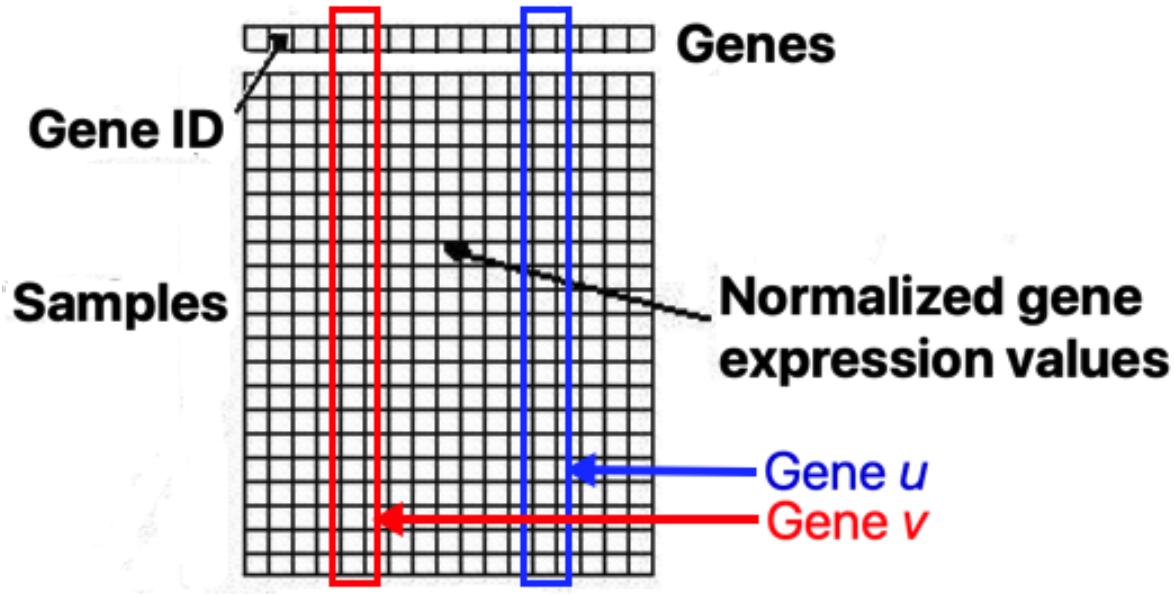
Count matrix X with shortlisted genes as columns (red and blue column represent genes u and v), samples as rows, and normalized gene expression as values

As a preprocessing step, genes *u* that do not satisfy a prespecified cut-off of mean expression and M-value across all samples [***X***_0,*u*_, ***X***_1,*u*_,…, ***X***_*S*–1, *u*_] are eliminated. The M-value is a measure of variability in gene expression based on the premise that the expression ratio of two stably expressed genes should be similar in all samples [29]. In other words, given expression matrix ***X*** and two stable genes *u*_1_, *u*_2_ in samples *j*_1_, *j*_2_, will follow 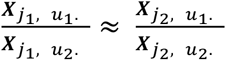. Based on this, the M-value is defined in the following two steps. First, the pairwise variation between gene *u*_1_ and gene *u*_2_ is calculated as the standard deviation of the log-fold changes between their expression levels across all samples:

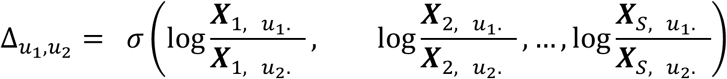

Second, the M-value of gene *u*_1_, is calculated as:

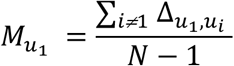

Subsequently, each remaining element in ***X*** is normalized by *max*(*X*).

### 2.2. Creation of perturbed and control networks

Given RNA-seq gene expression data from two biological conditions, CoVar first creates networks independently for each condition. Expression networks for the perturbed and control expression data are created using GENIE3 feature selection [27]. Feature selection techniques, such as logistic regression and extra-trees classification, select a subset of dimensions (or features) that contribute towards the prediction of an observed variable (called the label). Feature selection approaches often eliminate features with a low variation as they are non-informative with respect to the labels. Selection of a subset of informative features makes these techniques different from dimension reduction approaches (e.g., principal component analysis) or compression (e.g., using information theory) that alter the original representation of the variables [30, 31].

Given a pair of genes *u* and *v* (s.t. 0 < *u, v* < *N*), the purpose of applying network inference on gene expression data (***X***) is to predict how *u* influences *v*(*u* ≠ *v*), and vice-versa. GENIE3 calculates the influence of each gene on another, and the influence of gene *u* on gene *v* is represented as a directed weighted network with *N* genes as nodes. Note that GENIE3 networks are fully connected network with a directed link from each node to all other nodes.

Given the expression matrices for perturbed and control experiments, CoVar calculates the weighted directed genie matrices (Figure 2) for each network, where each directed link (*u, v*) has the weight *W*_(*uv*)_ ∈ [0,1], denoting the strength of the influence on the expression of node *v* by *u*. GENIE3 assumes that the expression profile of gene *u*(*x_u_*) is a function *f* of the expression of all genes *v*(*u* ≠ *v*) plus random noise *ϵ*, i.e.,

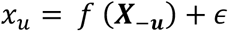

Here ***X***_−*u*_ is the vector containing the measurement of all vectors except *x_u_*, i.e., [*x*_0_, *x*_1_,…, *x*_*u*–l_, *x*_*u*+1_,…, *x_n_*].

**Figure 2:**
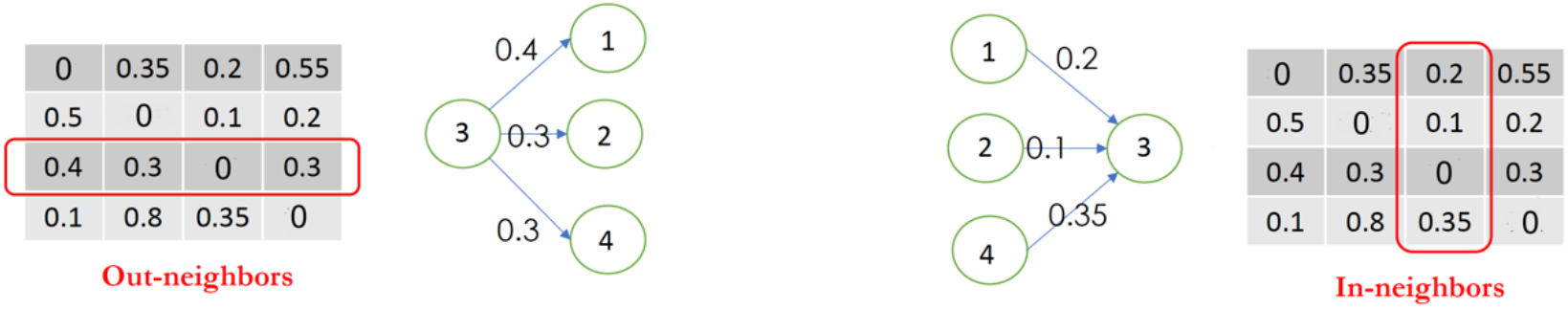
Out- and in-neighbor of gene 3 identified based on weights learnt from GENIE3

For each gene *u*, GENIE3 employs machine learning feature ranking to find the parameters *w* to function *f* that minimizes the error (*x_u_* – *f*(***X***_−*u*_))^2^.

### 2.3. Identification of variational and differentially expressed genes

CoVar next identifies variational genes whose neighborhoods in perturbed and control networks vary the most. Variational genes are not the same as differentially expressed genes. For each gene *u*, there is an out-degree GENIE3 weight vector [*w*_(*u*,1)_, *w*_(*u*, 2)_,…, *w*_(*u,N*)_] and in-degree weight vector [*w*_(1,*u*)_, *w*_(2,*u*)_,…, *w*_(*N,u*)_], respectively. Variational genes are defined as those exhibiting the highest variation in the in- and out-neighbor weights between the control and perturbed datasets. To gauge variationality, CoVar creates concatenated (in- and out-neighbor) weight vectors in both control and perturbed (*V^control^*(*u*) and *V^perturb^*(*u*)) as shown below.

Figure 2 shows in red the in- and out-neighborhoods of gene 3. The concatenated vector will be [0.2, 0.1, 0, 0.35, 0.4, 0.3, 0, 0.3].

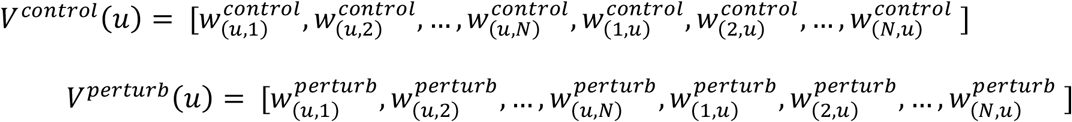

The variationality of gene *u* is the mean squared error of *V^contro1^*(*u*) and *V^perturb^*(*u*), i. e., *Mean Squared Error*(*V^contro1^*(*u*), *V^perturb^*(*u*))).

We contrast the variational genes with differentially expressed genes determined independently of changes in the expression of other genes. A differentially expressed gene (DEG) is one whose observed change in read count or expression value across two experimental conditions is statistically significant. For our analysis, we use the *DESeq2* [32] differential expression package and consider genes with an adjusted p-value < 0.05 to be differential.

### 2.4. Constructing the nearest neighbor network

Nearest neighbor (NN) is a machine learning classification technique in which a data point is assigned a class based on the class membership of the majority of its most similar (or nearest) data points [33]. CoVar employs NN to identify genes with the strongest influences on or are strongly influenced by the variational genes and constructs the nearest neighbor network constituting the variational genes, their neighbors, and their shared directed links.

CoVar assigns equal importance to all the variational genes. This approach greedily selects the neighbors with the highest incoming and outgoing link weights across all the variational genes. It initializes the NN network with variational genes of the perturbed network, iteratively adds the nodes with the highest incoming or outgoing link weight to the existing NN network and stops when there are links in the NN network. While this greedy approach is used to carry out the experiments, we present another NN construction approach (refer to Supplementary Method). Note that, unlike the GENIE3 perturbed and control networks, the NN network is not fully connected. It only possesses edges with the highest link weights among the variational genes and their neighbors.

### 2.5. Identification of modules through community detection

Modules or communities within networks can be identified through a measure called modularity. Networks with high modularity have dense connections between the nodes within modules, but sparse connections between nodes across modules [34]. Modularity in a directed network is calculated as:

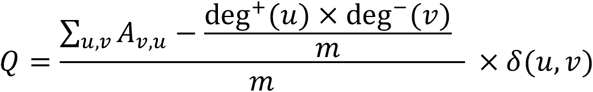

In the above equation, *A_v,u_* is 1 if (*u, v*) ∈ *G* and *δ* (*u, v*) = 1 if *u* and *v* belong to the same community, and 0 otherwise. We employ Louvain-Newman community detection for directed networks [35], which assigns community labels to nodes such that the modularity *Q* is maximized. It finds divisions within the network such that there are more directed links within communities than expected by chance [36]. Community detection was performed on the nearest neighbor network defined above (see 2.4) to determine modules.

### 2.6. Core network formation

The *k-core* of an undirected graph is the subgraph remaining where all nodes with degree less than *k* are removed. All the nodes in the *k*-core have a degree at least equal to *k* and the *k* + 1-th core is always a subgraph of the *k*-th core [37]. Core genes identified by CoVar possess two properties, namely *coordination* and *reachability*. Coordination is measured in terms of graph density, defined as the ratio between the number of edges and the maximum number of edges in a directed core graph *G* (*V, E*). It is calculated as:

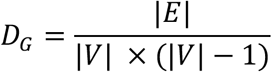

The reachability of a directed core network *G* (*V, E*), *R_G_*, is the fraction of total nodes in the overall network that have a path from at least one core node in *u* ∈ *V*. We employ the *undirected* core detection, where each core gene *u* has degree *deg*(*u*) = *k*, i.e., connections with *k* other core nodes. There is also a *directed* core variant of CoVar (refer to Supplementary Method), which preserves the directionality of the genes in the nearest neighbor network.

### 2.7. Combining variational, nearest neighbors, modules, and core genes across multiple runs

There may be slight variation in the variational and core genes in independent runs of the CoVar approach. This is because GENIE operates on randomized trees and the directed network it returns may vary slightly in edge weights across runs. We address this variability by combining the nearest neighbor networks across multiple runs to generate an integrated nearest neighbor network, by applying the following three-step process:

1. The integrated network is initialized as an empty network and populated with all the nodes and edges appearing in all the runs.
2. The edges that appear in less than a prespecified integral parameter *ρ* (where 0 ≤ *ρ* ≤ number of runs) are removed from the integrated network. The isolated nodes, i.e., nodes that are not connected to any other nodes in the trimmed (integrated) network are removed from it.
3. The variational, nearest neighbor, and core genes in the combined network are determined as described above (2.3 – 2.6).

For all experiments performed in this study, we used the following parameter settings.

### 2.8 Small world networks

Small world networks are networks in which most nodes are not neighbors of one another, and any node is reachable from any other node in a few hops (edges) [38]. The Barabási Albert preferential attachment growth model is a small world network generation approach in which we start with a few densely connected nodes, and subsequently, nodes are iteratively introduced into the network *G* (*V,E*), where *V* is a finite, non-empty set of objects called vertices (or nodes); and *E* is a (possibly empty) set of 2-vertex subsets of *V*, called edges [39]. Any new node *v* prefers to be attached to a well-connected node *u* with probability given by:

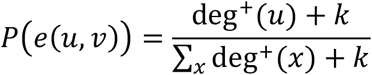

Here *k* is the small world parameter that assumes integral values and controls the overall connectivity of *G* (*V, E*). Specifically, a low value of *k* results in networks where very few nodes are highly connectivity (shown in red in Figure 3a); conversely, higher *k* generates networks where most nodes have average connectivity (Figure 3b). We employ Eq. 1 to generate networks with low and high *k* values that vary greatly in network properties.

**Figure 3:**
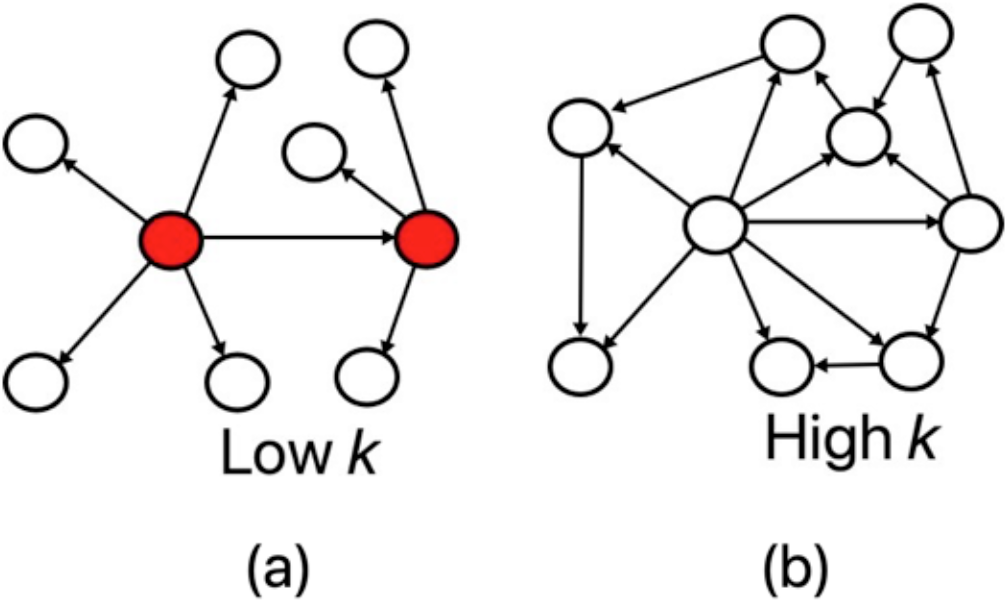
Small world networks. Networks with (a) a low value of the small world parameter k results in few highly connected nodes (shown in red); and (b) a high value of k creates networks with most nodes having similar connectivity.

### 2.9. Differential coexpression analysis

Differentially coexpressed genes (DCEGs) are groups of genes whose coexpression varies across conditions. DCEGs are coexpressed in the control and not in the perturbed expression datasets, or vice versa. CoXpress [40] is a differential coexpression analysis package, which applies hierarchical clustering to find coexpressed genes in one experiment and then uses a resampling technique to determine whether the same group of genes is coexpressed in experiments.

### 2.10. Yeast validation dataset

We validate CoVar using gene expression data from a study on mitochondrial genome depletion in yeast (GEO accession: https://www.ncbi.nlm.nih.gov/geo/query/acc.cgi?acc=GSE162197) [41]. We consider control and perturbed datasets of a wild-type and a mutant strain in which the mitochondrial genome has been depleted. The *control samples* are EV1 mRNA-Seq WT replicates 1-3) GSM4946300, GSM494630, GSM4946302), while the *perturbed samples* are EV1 mRNA-Seq r0 replicates 1-3 (GSM4946315, GSM4946316, GSM4946317).

## 3. Results

### 3.1. Overview of CoVar

CoVar utilizes machine learning and complex network theory to identify variational and core (or central) genes from gene expression data that are altered across two different biological conditions. Expression data from perturbed and control conditions are each represented as networks where nodes are genes and expression relationships are directed edges (*u, v*) with weight *w_u, v_* ∈ [0,1] that represents the strength of the influence of gene *u* on the expression of gene *v* (Figure 4a; Methods 2.2). Genes that show the largest changes in the strength of influence relationships (weights) with neighbor genes are identified as variational genes (Figure 4b; Methods 2.3), providing an initial set of genes highly affected by the biological perturbation. Genes with strong connections to the variational genes denoted as their nearest neighbors (Figure 4c; Methods 2.4), are determined and together with the variational genes form a nearest neighbor network. Modules, or network communities, are identified to focus on well-connected groups of genes (Figure 4d; Methods 2.5). Within each module, core genes are determined that have two specific properties: (1) *coordination*, meaning they form a densely interconnected core network with other core genes; and (2) *reachability*, meaning they have directed paths to the bulk of the non-core genes, influencing the latter’s expression (Methods 2.6). As the initial perturbed and control networks are determined using a non-deterministic machine-learning algorithm, this process is repeated multiple iterations to find a final set of genes consistently identified variational, nearest neighbor, and core genes (Methods 2.7).

**Figure 4:**
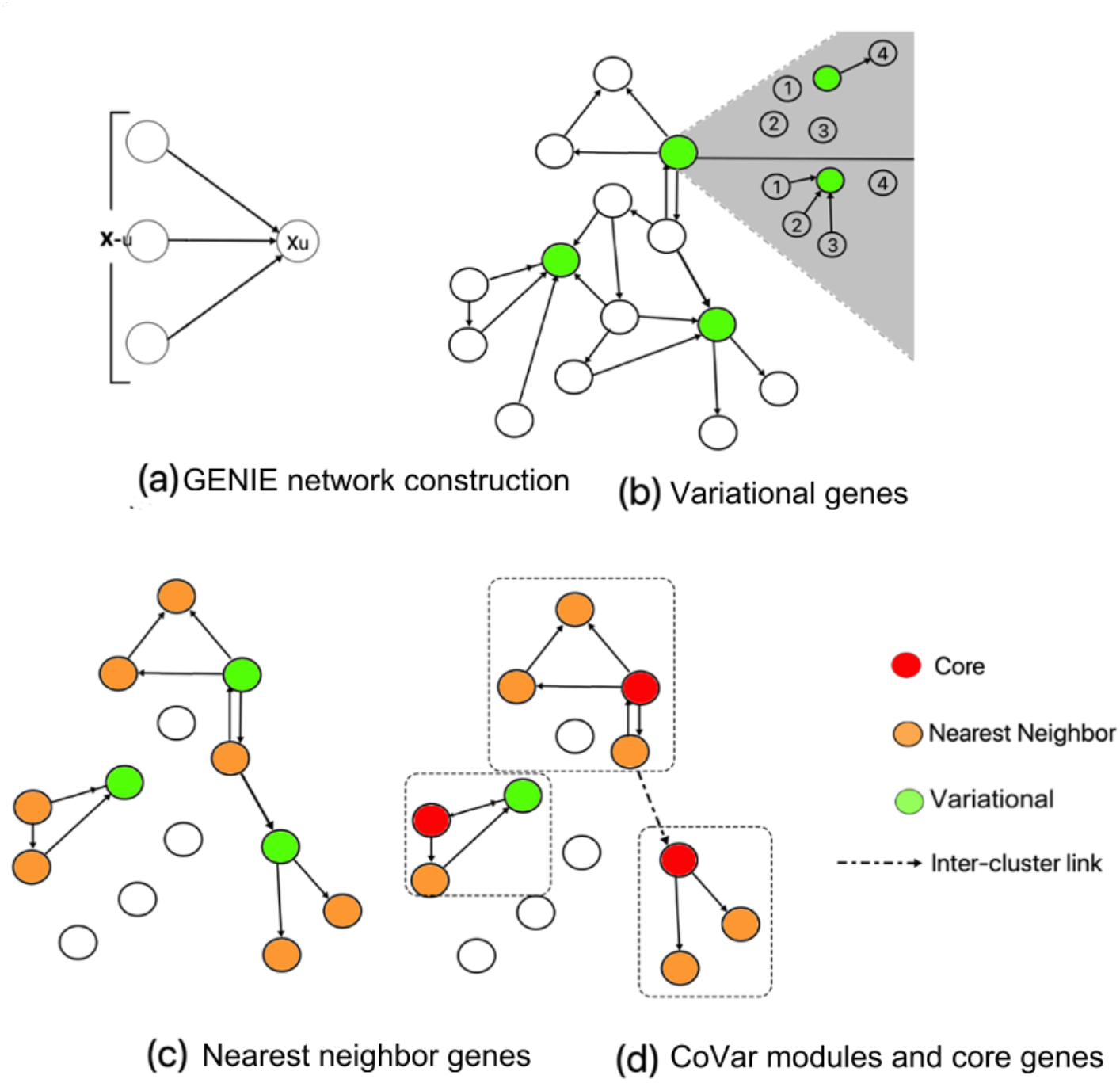
Steps in the CoVar method. (a) GENIE uses randomized trees defines the edges and weights connecting genes in the two separate conditions; (b) variational genes (colored green) are identified based on differences in edges across the control and perturbed networks; (c) nearest neighbors (colored orange) for the variational genes are identified in the perturbed network; (d) modules are defined and core genes are identified for the nearest neighbor networks in each module.

### 3.2. Characterization of Variational Genes

CoVar measures the variationality of genes between the control and perturbed datasets in terms of the mean squared error of the edge weights between the corresponding CoVar networks. In this experiment, we study whether these variational genes also exhibit a change in the mean (and standard deviation of expression values in the two datasets. To this end, we synthetically generated expression profiles of 16 genes that varied in two ways: (1) change in mean expression between the control and perturbed networks and (2) variance in gene expression profiles. Table 2 shows the range of the gene expression values for low and high variance as well as the change in mean expression across control and perturbed data. We applied the CoVar pipeline to generate the control and perturbed networks, where the expression values of the genes lie in the range specified in Table 2. Following this, we calculated the variational expression of the 16 genes measured in terms of mean squared error or MSE.

**Table 1:** Parameter settings used in CoVar evaluations.

| Parameter | Value |
| --- | --- |
| M-value cut-off | 0.32 |
| Mean expression cut-off | 60 |
| Number of perturbed and control samples | 3, 3 |
| Number of trees in feature selection | 10000 |
| Percentile cut-off for variational genes | 97.5 |
| Percentile cut-off for nearest neighbor links ($Z$) | 99.95 |
| Number of runs | 25 |
| Nearest neighbor approach | Greedy |
| Core network approach | Directed |

**Table 2:** Range of the gene expression values for 16 genes having low and high variance as well as change in gene expression across control and perturbed data.

| Gene ID | Expression range in control | Expression range in perturbed |
| --- | --- | --- |
| Low var. and low change<br>(4 genes) | [10, 20], [10, 20],<br>[10, 20], [10, 20] | [15, 25], [5, 10],<br>[15, 25], [5, 10] |
| High var. and low change<br>(4 genes) | [10, 50], [10, 50],<br>[15, 55], [5, 45] | [15, 55], [5, 45],<br>[15, 55], [5, 45] |
| Low var. and high change<br>(4 genes) | [10, 20], [190, 195],<br>[200, 210], [10, 20] | [200, 210], [10, 20],<br>[200, 210], [10, 20] |
| High var. and high change<br>(4 genes) | [10, 50], [150, 200],<br>[10, 50], [150, 200] | [200, 250], [10, 50],<br>[200, 250], [10, 50] |

We recorded the average MSE from 10 iterations of the experiment, where the variational expression is calculated based on the variability of only in-neighbors (Figure 5a) and both in- and out-neighbors (Figure 5b) across the control and perturbed networks. These results show that as might be expected, genes with low variance and large changes in mean expression across conditions, characteristics of significantly differentially expressed genes, were the top genes in terms of variability in strengths of relationships with neighbor genes. In addition, though, we see that genes with high variance across conditions and large changes in mean expression also resulted in higher MSE. Together, these suggest that variational genes are likely to be differentially expressed as well but maybe different from the most significantly differentially expressed, which are often penalized for high variance. And even genes with low changes in expression may result in high MSE, though less frequently than genes with larger changes. Thus, variational genes identified by CoVar are not simply the most highly differentially expressed genes.

**Figure 5:**
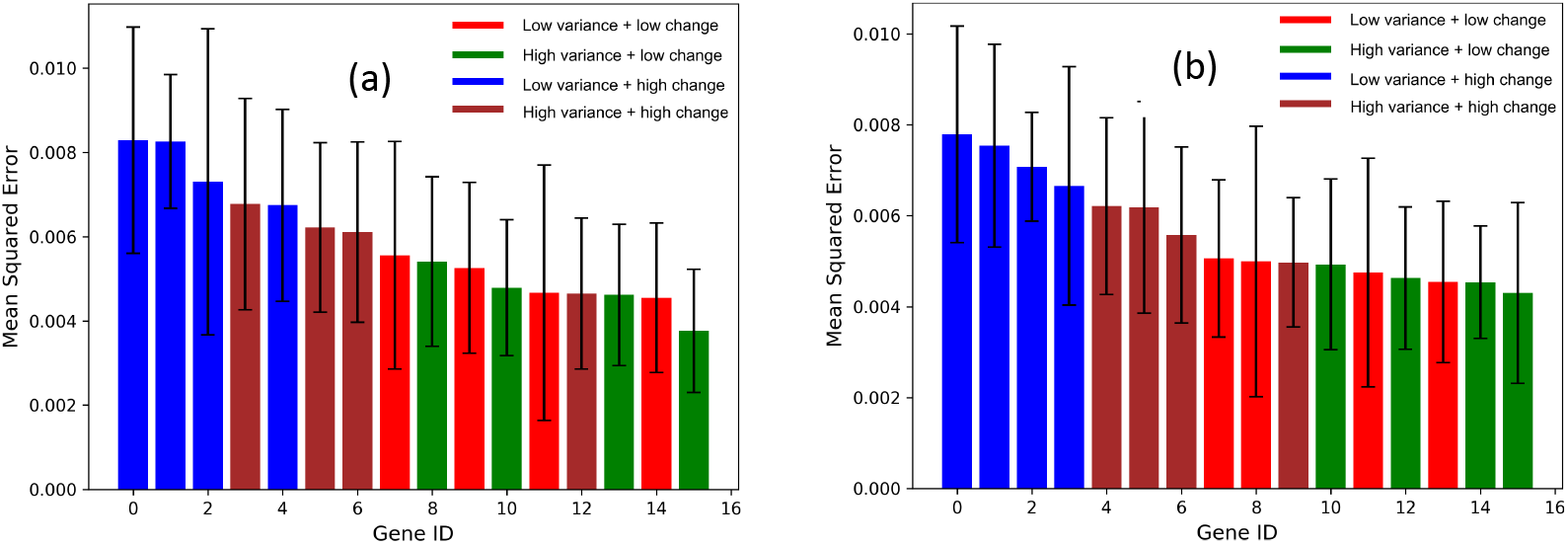
Genes with low and high variance and change in mean expression across control and perturbed data sorted in the decreasing order of their mean squared error

### 3.3. Characterization of Core Genes

As stated previously, we want to identify core genes that are both highly coordinated amongst themselves while also having high reachability to other genes within the network. To determine how well we capture these characteristics, we created 1000 small world networks with small world parameters *k* = 3 and *N* = 100,200, · · ·, 500 nodes each and determined *c* core nodes in each network. We also randomly chose *c* core nodes and summarized the number of times that the coordination and reachability (Methods 2.6-2.7) of the randomly chosen nodes equals or exceeds those of the CoVar core nodes in the corresponding nearest neighbor network. Table 3 shows that the CoVar core nodes offer the best mix of coordination and reachability – while no randomly chosen node set has higher coordination than the core nodes, only a fraction (0.121 — 0.415) of these sampled sets equal (but do not exceed) the reachability of the CoVar core nodes.

**Table 3:** Number of times the coordination and reachability of randomly selected nodes in 1000 small world networks of *k* = 3 equals or exceeds those of the code nodes.

| No. of nodes ($N$) | Coordination ($>, \geq$) | Reachability ($>, \geq$) |
| --- | --- | --- |
| 100 | (0, 0) | (0, 121) |
| 200 | (0, 0) | (0, 145) |
| 300 | (0, 0) | (0, 329) |
| 400 | (0, 0) | (0, 395) |
| 500 | (0, 0) | (0, 415) |

We were interested in how the distribution of the degree of node edges affects core gene identification. As described in the Methods (section 2.8), the small world property of networks can be controlled using the parameter *k*. A small *k* generates a network with a few highly interconnected hub nodes that are connected to the majority of the non-hub nodes. Increasing *k* results in a network of uniform connectivity and fewer hubs. We found that in all cases, the reachability of the core nodes remained very high, but that the fraction of nodes of identified core nodes increased as *k* increased (Figure 6a). Interestingly, the coordination was similar across all runs except for the one with the highest *k*, where coordination was substantially increased.

**Figure 6:**
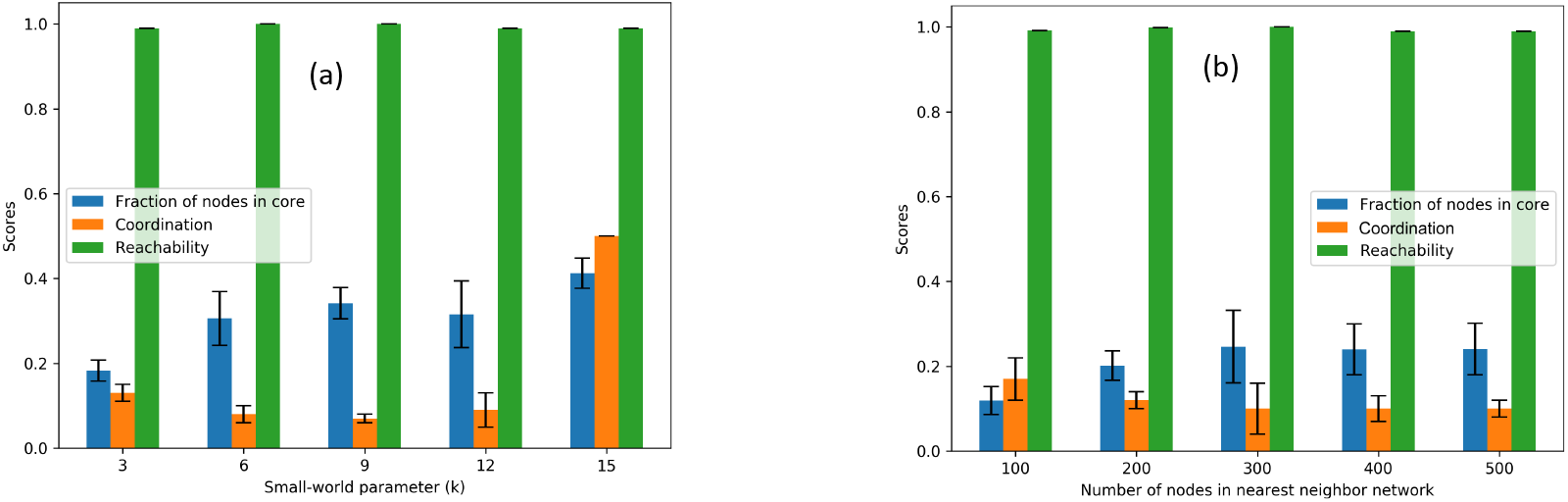
Coordination and reachability of small world networks for varied (a) smallworld parameter (*k*) and number of nodes *N* = 200 and (b) number of nodes and *k* = 3.

We also wanted to determine whether the number of nodes in the nearest neighbor network had a significant effect on core gene identification. We varied the number of nodes *N* while keeping *k* consistent. Again, the reachability remained high across all settings for *N*, while the fraction of nodes identified as core nodes was inversely correlated with the coordination (Figure 6b). As *N* increased, the fraction of nodes in the core also increased while the coordination decreased. This shows that these basic properties of coordination and reachability of the networks are stable across network sizes.

### 3.4. Analysis of gene networks altered by mitochondrial genome depletion in yeast

To demonstrate the ability of CoVar to identify genes that contribute to altered expression in a perturbed state, we analyzed expression data from yeast generated under two conditions: a normal state and a perturbed state in which the mitochondrial genome was deleted in a study by Lui et al. [41]. They discuss that mitochondrial dysfunction and oxidative phosphorylation (OXPHOS) impairment due to mitochondrial genome depletion leads to adaptive responses, where the downregulated processes are associated with ribosome biogenesis and the upregulated processes are mitochondrial translation, mitochondrial gene expression, and carbohydrate metabolic process. Upon deletion of the mitochondrial genome, a primary consequence is a reduction in mitochondrial membrane potential.

We filtered non-coding genes and lowly expressed genes (mean expression < 50 or M-value < 0.32) resulting in 1843 genes remaining. We carried out 25 independent runs of CoVar on 3 control and 3 perturbed samples, each of which generates 1843-node control and perturbed networks with slightly varying edge weights. For each run, the top 47 variational genes (97.5 percentile highest mean squared error (MSE) were selected. We note that 22 out of the 25 most consistently identified variational genes across the 25 runs are also differentially expressed (p-adjusted cut-off 0.05). In each run, the nearest neighbors were then determined, and the nearest neighbor network was constructed with 547 genes on average. Within each nearest neighbor network, modules of highly connected genes were determined, with there being 3-4 modules consistently found across the 25 runs. Finally, core genes were identified within each module with an average of 63 core genes per run. Table 4 shows summary statistics of the core, nearest neighbor, and variational genes across these 25 runs, where the mean density and reachability of the core nodes in the perturbed network are 0.067 and 0.247 respectively. It is noteworthy that the mean MSE of the variational genes is nearly double that of the entire set of 1843 genes.

**Table 4:** Summary statistics of the core and variational genes across 25 runs

| **Metric** | **Mean** | **St. Dev.** |
| --- | --- | --- |
| Density of core genes | 0.420 | 0.117 |
| Reachability of core genes | 0.999 | 0.003 |
| (Mean) MSE of variational genes | 0.021 | 0.001 |
| (Mean) MSE of nearest neighbor network | 0.012 | 0.001 |
| Mean (and standard deviation of) the number of nearest neighbor genes across 25 runs | 547.52 | 23.73 |
| Mean (and standard deviation of) the number of core genes across 25 runs | 96.88 | 12.44 |
| Mean (and standard deviation of) the number of modules across 25 runs | 3.96 | 0.52 |

We compiled the data across the 25 independent runs to generate a final network of three distinct modules consisting of 267, 113, and 94 genes for a total of 474 genes (section 2.7; Supp Table 1 – the table of genes). Of these, 86 genes were designated as core genes, with 45 in module 1 (M1), 24 in module 2 (M2), and 17 in module 3 (M3). We examined connections in the final network for the core genes and found that on average, each core gene had 16.99 out edges and 17.76 in edges compared to 2.75 out edges and 2.58 in edges for non-core genes. This is expected based on the focus on reachability in defining core genes.

To determine whether the genes identified across the whole network corresponded to previously published results, we performed an enrichment analysis of the set of all nearest neighbor genes using Enrichr [42]. We found that among the top gene ontology terms are translational elongation, structural integrity of the ribosome, translation termination as well as post-transcriptional modification regulated by the untranslated region of the mRNA, and protein metabolism. Indeed, these results match well with the general cellular processes cited as being affected by mitochondrial genome depletion.

To better understand the genes contained within the modules, we performed a similar pathway enrichment analyses on genes in each of these independently. Again, we see that the top GO terms and pathways for all three modules were related to the ribosome or translation, and many of the remaining were related to mitochondrial function and metabolic processes (Supp Table 1). As the three modules have mutually exclusive sets of genes, we assessed whether there were distinct functions associated with each module. In the first module (M1), we find enriched gene sets related to translation very generally, along with mitochondrial organization, protein degradation, and protein transport into multiple nuclear locations including the mitochondria and endoplasmic reticulum. This module includes the enzyme Mitochondrial intermembrane space Import and Assembly protein 40 (*MIA40*) gene, a key component of the MIA pathway that imports proteins to the inner mitochondria and facilitates their correct oxidative folding [43]. Module M1 also includes 20 transposable element genes. Increased transposition activity has been associated with response to stress conditions in yeast and other organisms [44]. In the second module (M2), we see distinct RNA processing functions, especially for ribosomal RNAs, but also related to respiration that include several cytochrome c oxidase (COX) genes and isocitrate dehydrogenase 2 (*IDH2*). The third module (M3) is specifically enriched for ribosomal genes involved in ribosome assembly as well as key genes in lipid, sterol, and fatty acid metabolism such as the mitochondrial NADH-cytochrome b5 reductase *MCR1*, the farnesyl diphosphate synthase *ERG20* (*FPP1*), and the C-5 sterol desaturase *ERG3*. Thus, we find that the identified modules provide important information on relationships between genes involved in more specific cellular functions than previously noted.

### 3.5 Comparison of expression change characteristics between differentially expressed genes and variational genes

Identification of key genes driving a perturbed molecular state is most often done by determining genes that are significantly differentially expressed. These analyses focus on each gene independently without considering how a gene may affect or be affected by other genes. We wanted to understand how variational genes identified by CoVar compared with differentially expressed genes. We identified differentially expressed genes (DEGs) using DESeq2 (section 2.3) and ranked genes based on their computed adjusted p-value. We considered genes with adjusted p-values < 0.05 to be differentially expressed.

We hypothesized that variational genes differ from the differentially expressed genes (DEGs) in how they are altered in relation to genes that are correlated in expression, such as neighbor genes in a network, as opposed to simply their own change in expression. Consider a DEG and a variational gene, *u_d_* and *u_v_*, with expression vectors denoted as ***x_u_d__*** and ***x_u_v__***. A PCA analysis using these vectors would focus primarily on the change in expression of the single gene. Figure 7a shows the first two components (PC1 and PC2) from a hypothetical principal components analysis on ***x_u_d__*** and ***x_u_v__*** in the control and perturbed expression data samples, we denote hereafter by 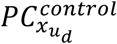, 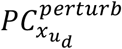, 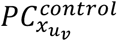, 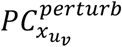. We intuit that there will a greater Euclidean distance between 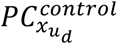 and 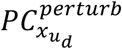 for the DEG than between 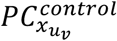 and 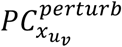 for the CoVar variational gene. This is because the DEGs undergo a change in their *own* expression profiles of the control dataset with respect to their perturbed dataset.

**Figure 7:**
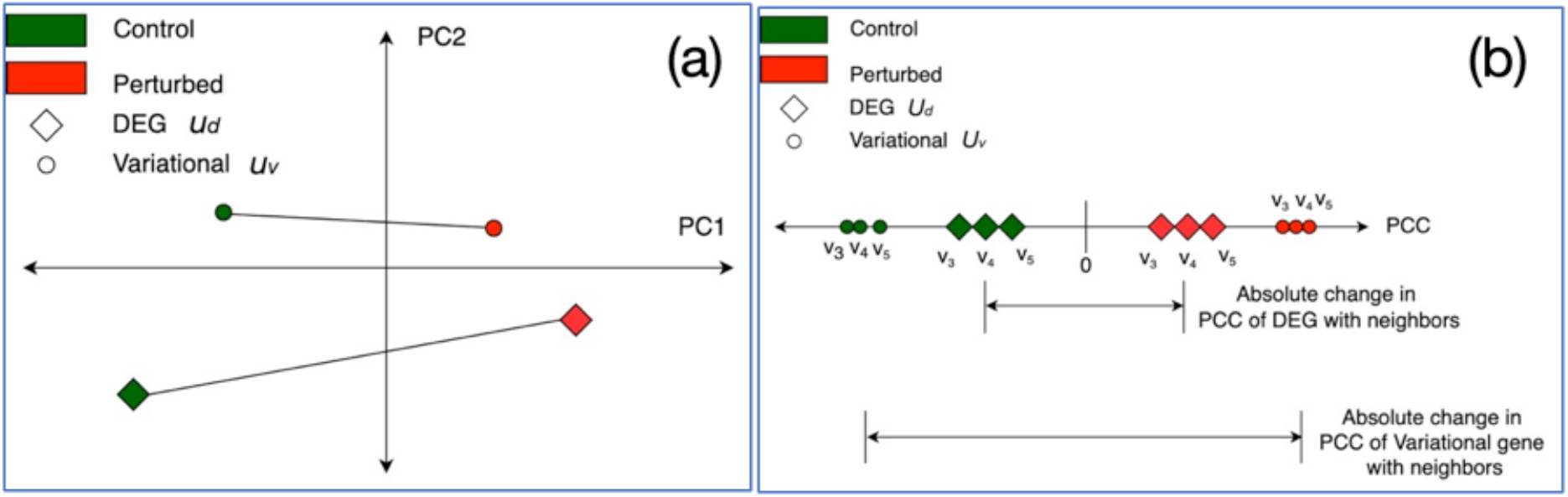
Proposed model of differences between variational and differentially expressed genes in control versus perturbed expression data. (a) Principal component analysis of expression vectors of DEG and variational gene *u_d_* and *u_v_*, where we expect differentially expressed genes individually show greater change across conditions than variational genes; (b) Mean change in correlation between expression of *u_d_, u_v_* and the neighbor genes, where we expect that variational genes will exhibit a greater change in this correlation.

In contrast, the less significantly differential gene *u_v_* may be identified as a CoVar variational gene when its variationality emanates from a greater relative change in expression with respect to its neighbors across the control and perturbed data. This change is partly measured by the mean absolute change in the Pearson correlation coefficient (PCC) in the expression of *u_v_* and its neighbor genes between the control and perturbed samples. We illustrate this with an example of three other neighbor genes *v*_3_, *v*_4_, *v*_5_ in Figure7b. We can quantify this relative change in the neighborhood of a gene across the two datasets as 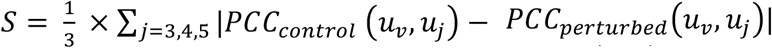. Thus, measure variationality as a function of the relative change in expression between a given gene and other genes, caused by the combination of differential expression of the gene, its neighbors, or both. In this hypothetical example, *u_v_* has greater variationality than *u_d_* even though it is not as significantly differentially expressed (Fig.7b).

To test this hypothesis on the yeast dataset, we carried out the above principal component analysis (PCA)- and Pearson correlation coefficient (PCC)-based study. Specifically, for any given gene, we first applied PCA and recorded the Euclidean distance between the vectors comprising the first two components (PC1 and PC2) in the control and perturbed networks. Figures 8a and 8b depict these distances for all of the DEGs and variational genes. We see that the DEG have a higher mean distance than variational genes indicating DEGs undergo a greater change in their own expression vector between the perturbed samples and the control samples, similar to our hypothetical example. Second, we calculated the PCC for each of the DEGs and all other genes and similarly for variational genes and all other genes in one of our 25 CoVar iterations. Here, we find that the mean change in the PCC of the expression vector of the variational genes with all other genes between control and perturbed data is higher than the mean change for DEGs (0.907 vs 0.838. Figure 8c and 8d), similar to our hypothetical example. Note that the average and standard deviation in correlation change (*S*) of the variational genes across 25 runs are 0.8477 and 0.269, respectively.

**Figure 8:**
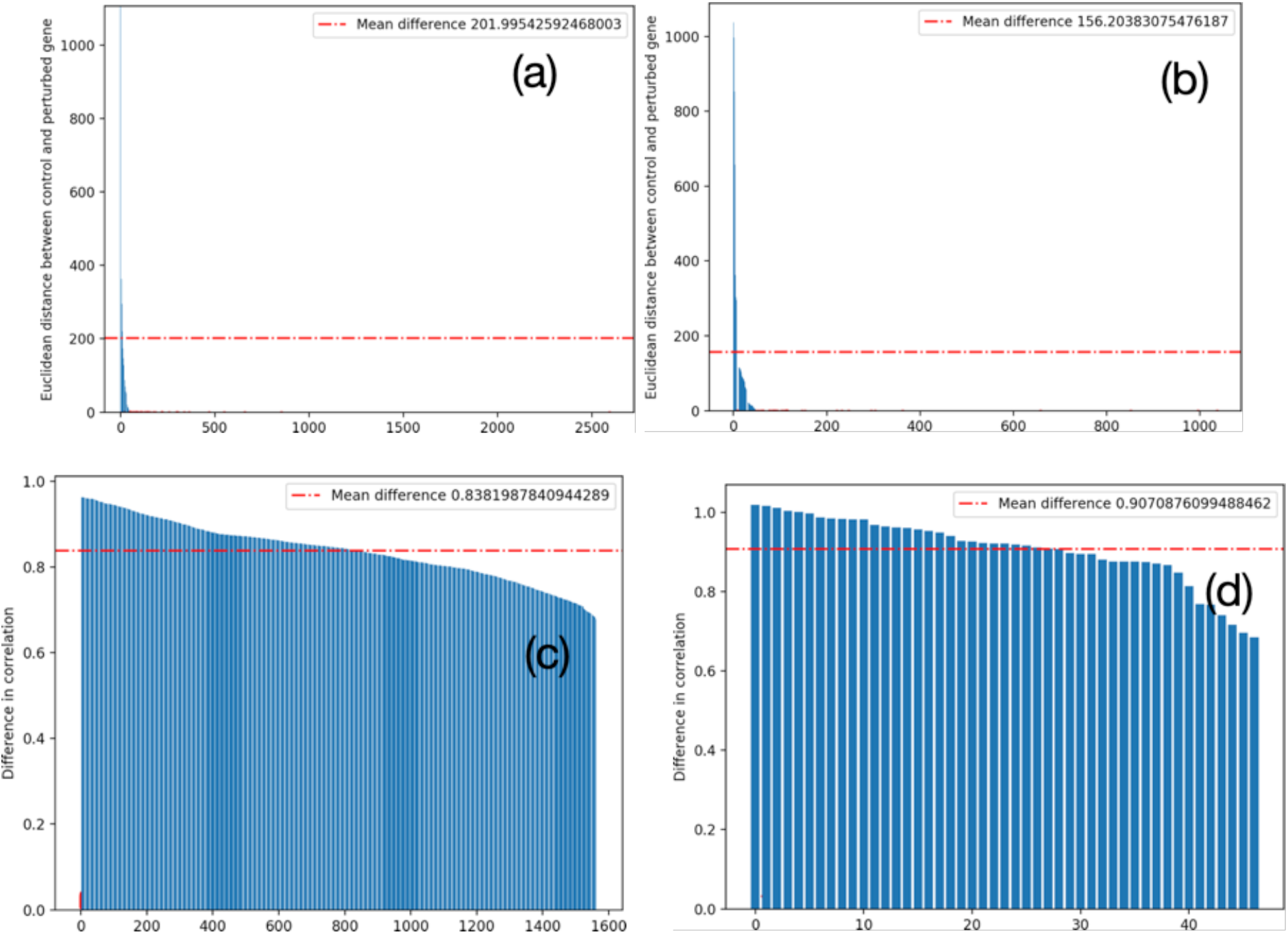
Differentially expressed genes (DEGs) versus variational genes. Principal component analysis of expression vectors of (a) DEGs (b) variational genes; Mean change in Pearson correlation between the expression of (c) DEGs and (d) variationalgenes against other genes

We performed a direct comparison of relative change in the neighborhoods of the DEGs and variational genes in terms of the correlation change C. The variationality of a gene is measured in terms of the MSE of its control and perturbed weight vectors of the edges with its neighbors in the GENIE networks (section 2.2). The goal here is to contrast the C scores of the DEGs (genes likely to have a high absolute value of log2FC for a low adjusted value) and variational genes across the 25 runs (with higher mean MSE). Figures 9a and 9b show that the MSE of the DEGs is uncorrelated with both the absolute log2FC (*PCC* = −0.088) and adjusted p values (*PCC* = −0.064), while Figures 9c shows that the correlation change of the variational has a low but positive association (PCC 0.106) with their MSE. This supports that the variational genes exhibit a greater overall change in expression relative to the other genes than DEGs.

**Figure 9:**
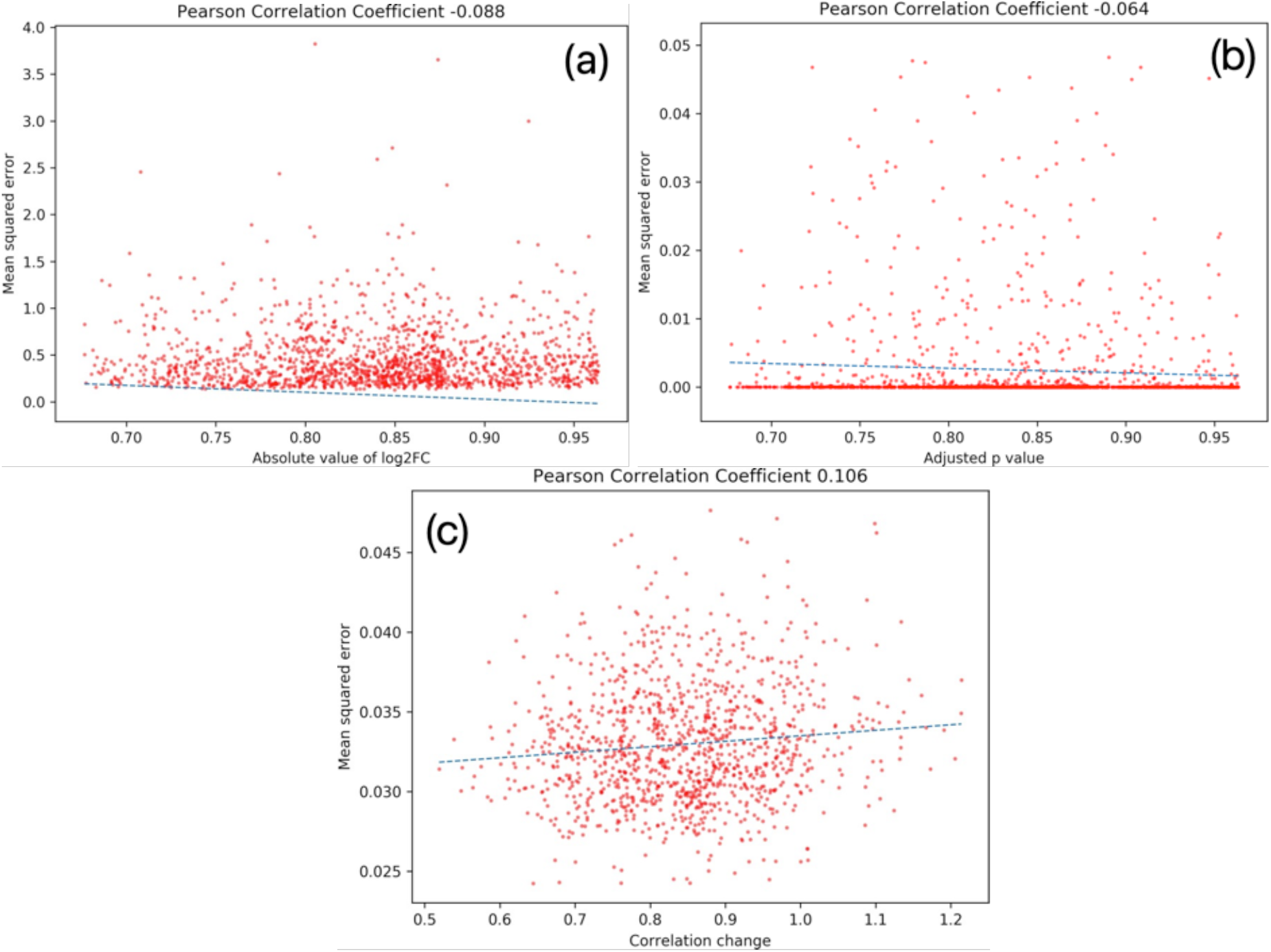
Pearson correlation between mean squared error (between the control and perturbed networks) and (a) absolute value of log fold change and (b) adjusted p value of the differentially expressed genes, as well as (c) correlation change of variational genes in the 25 runs

### 3.6 Comparison of cellular functions between differentially expressed genes and core genes

We further compared our core genes more directly with the most differentially expressed genes to see how their reported biological functions aligned with the change in experimental conditions. First, we ranked our core genes in decreasing order of their frequency of occurrence as a core gene in the 25 runs of CoVar. We also ranked the DEG genes based on their adjusted p-value to represent the significance in their differential expression. We find that 69 of the 86 cores genes were differentially expressed (adjP < 0.05; Supp Table S1), indicating that 17 (20%) were not. Further, the average DEG ranking for core genes was 234.19. These show that while most core genes were differentially expressed, they were not simply the most differential genes.

As mentioned earlier, most of the core genes with a frequency of occurrence of at least 10 (74 out of 86) are DEGs and are amongst the most frequent variational genes. The top two ranked DEGs, Isocitrate DeHydrogenase 2 (IDH2; upregulated on mitochondrial genome loss) and Vacuolar Transporter Chaperone 3 (VTC3; downregulated), were also core genes (ranked 27^th^ and 28^th^). IDH2 is a mitochondrial enzyme that is critical for a properly functioning oxidative phosphorylation system [45]. VTC3 is one of four subunits of the vascular chain transporter that helps control polyphosphate levels, critical for energy metabolism [44]. Other than these, though, only one other top-40 ranked DEG was identified as a core gene in the final combined model. We note that other top-ranked DEGs did appear as a core gene in at least one of the CoVar iterations. For example, Heat Shock Protein 12 (HSP12), associated with heat shock, osmostress and oxidative stress [46], ranked 3rd among DEGs and appeared in 4 runs as core genes. ARGinine requiring 3 (ARG3) is the 5th ranked DEG and a core gene in 7 runs that has been shown to be induced by heat shock and by other cellular stress inducers [47]. Finally, Multiprotein Bridging Factor 1 (MBF1), ranked 9th among DEGs and a core gene in 5 runs [48], is associated with developmental processes and abiotic stress, particularly heat stress, in plants.

Many notable core genes were not among the top ranked DEGs or not DEGs at all. The Blocked Early in Transport 1 (BET1) gene was the most consistently chosen core gene across the 25 iterations but was only the 374th most significant DEG (Supp Table S1). It had the largest number of edges in the final network (21 in edges, 75 out edges). BET1 is a membrane protein involved in vesicle transport between the endoplasmic reticulum (ER) and the Golgi apparatus that is upregulated on mitochondrial genome loss. The orthologous Bet1 protein in Drosophila was determined to be necessary for the maintenance of the mitochondrial genome, with prolonged knockdown of Bet1 associated with an increase in mitochondrial membrane potential [49]. In this study, expression of BET1 is increased upon mitochondrial genome loss, presumably in an attempt to maintain a normal membrane potential. The core gene Copper TRansporter 2 (CTR2) gene (35^th^ ranked core gene), a low-affinity copper transporter that is thought to make available copper stored in vacuoles, was not differentially expressed (adjP = 0.13, Supp Table S1). Interestingly, the related gene Copper TRansporter 1 (CTR1), a high-affinity copper transporter and part of the final nearest neighbor network but not a core gene, was significantly upregulated. Copper is essential to several mitochondrial functions including the electron transport chain and energy metabolism. Copper deficiency has been linked to increased mitochondrial membrane potential [50]. Together, these suggest that maintenance of the membrane potential upon mitochondrial genome loss may depend on access to intracellularly stored copper and that this process is driven by CTR2, even though its relative expression level is not greatly changed.

Other notable core genes include Prohibition Complex subunit 1 (PHB1; 4^th^ ranked core gene, 176^th^ ranked DEG, upregulated), which contributes towards mitochondrial homeostasis and differentiated T cell survival [51]; Mitochondrial Ribosomal Proteins of the Large subunit MRPL20 and MRPL44 (29^th^ and 71^st^ ranked core genes, 87^th^ and 109^th^ ranked DEGs, upregulated) that are critical for protein synthesis within the mitochondrion [52]; OLEic acid requiring 1 (OLE1; 37^th^ ranked core gene, 51^st^ ranked DEG, upregulated), which is involved in mitochondria inheritance [53]; and Proteinase C 1 (PRC1; 16^th^ ranked core gene, 207^th^ ranked DEG, upregulated) that regulates mitochondrial biogenesis during erythroid development [54].

### 3.7 Comparison of CoVar modules with differentially coexpressed genes

We examined the validity of the coexpression of clusters of genes identified by CoVar analysis by comparing them with genes identified by differential coexpression analysis (section 2.9). Recall that CoVar identifies modules comprising variational genes and their nearest neighbors in the perturbed network (section 2.4). Therefore, it is fair to compare the overlap between the genes in the CoVar modules and genes coexpressed in the perturbed dataset but not in the control dataset. We find that several sets of genes are both differentially coexpressed and belonging to the same CoVar modules. BET1, SPC3, and VOA1 form a differential coexpression set have a mean core frequency of 12.66 in 25 runs and appears in the same CoVar module (M1). Similarly, differentially coexpressed genes cPRC1, YNL284C-B, YPR158C-D, and AIM6 have a mean core frequency of 10.25 and appear in CoVar module M1. Also, differentially coexpressed genes DTD1 and SDH2 have a mean core frequency of 10 and are present in CoVar module M2. RPS0B, URM1, and RCR1 are differentially coexpressed, appear in CoVar module M3, and have a mean core frequency of 9.33.

## 4. Discussion

We present a network analysis framework, called *CoVar*, that analyzes control and perturbed expression data to identify central genes contributing to the perturbation. Given control and perturbed gene expression datasets, CoVar employs machine learning-based to create separate networks with directed links having weights commensurate with the influence of a gene on the expression of another. Unlike existing approaches that measure the importance of genes in terms of their own differential expression across control and perturbed datasets, CoVar identifies variational genes whose expression profiles change with respect to other genes. It selects central genes from the nearest neighbor network comprising the variational genes and their nearest neighbors in the perturbed network. These central genes exhibit both strong network ties with other central genes (termed coordination) as well as pathways to influence other nearest neighbor genes (reachability).

We demonstrate the efficacy of CoVar using synthetic and real expression datasets. First, we show that increased changes in mean expression across control and perturbed data samples, more so than high variation in expression in each dataset, are important for being identified as variational; but not all of the top variational genes are necessarily those with the largest change in expression. Second, we show that CoVar successfully identifies core genes that maximize coordination and reachability with other genes in the network, critical characteristics of genes with widespread influence. Finally, our analysis of wild-type and mitochondrial genome-depleted yeast datasets reveals that genes can be variational due to the combined effect of the differential expression of itself and its neighbor genes. A significant number (20%) of the core genes identified do not display significant differential expression between the two conditions, thus being among the most significantly differentially expressed does not necessarily translate to being a core gene. Yet importantly we find prior connections between these genes and the phenotype being studied. More broadly, we find that the variational and core genes identified by CoVar have functions and are involved in pathways that are unsurprisingly altered upon mitochondrial genome loss, not all of which were identified in a differential expression analysis of these data.

The CoVar approach is generalizable in many ways. First, while our choice of GENIE3 as the inference tool was guided by its capability of providing an exact measure of directed regulatory influence of each gene on the other, CoVar can operate on any input network generated through our means such as co-expression, Bayesian, Gaussian, or machine learning models. Moreover, the application of the existing network centrality measures can present new insights, as we have demonstrated during our analysis. Second, CoVar offers different modalities at each step. The choice of mean expression and M-value cut-off determines the size of the initial gene list and consequent variational and core genes. Variational genes can be alternatively estimated based on the in-degree and out-degree or both (section 3.2). Nearest neighbor identification has coverage and greedy approaches, and the core network can be directed or undirected. These modes can be selected to infer varying core genes based on the number of samples as well as the number or types of entities (i.e., genes, miRNAs) of the genomic data.

Third, CoVar is extensible to adapt to time-varying datasets. In addition to analyzing the variabilities between the control and perturbed networks, it can pinpoint the variations introduced within the control and perturbed networks over time. Fourth, there are several ways to incorporate prior knowledge about transcription, pathways, ontology, diseases/drugs, or cell types. For instance, one can generate nearest neighbor networks consisting of the known regulators of genes of interest or downstream genes of known transcription factors before the identification of the core genes. Moreover, one can find the paths in the control or perturbed networks, where the genes are connected by the highest edge weights. This approach lends itself to the analysis of other datatypes (such as chromatin, miRNA, protein expression), where the variabilities of the combined weights of the network paths can reveal central genes specific to the perturbation.

## Contributions

S.R., T.F. and S.S. conceptualized the work. S.R. performed data curation, developed methodology, performed analysis, and wrote the original draft. T.F. and S.S. validated the methodology and reviewed/edited the manuscript.

## References

[1] J. Churko, G. Mantalas, M. Snyder, and J. Wu. Overview of high throughput sequencing technologies to elucidate molecular pathways in cardiovascular diseases. Circulation research, 112(12):1613–1623, 2013.

[2] A. Aalto, L. Viitasaari, P. Ilmonen, L. Mombaerts, and J. Goncalves. Gene regulatory network inference from sparsely sampled noisy data. Nature communications, 11(1):1–9, 2020.

[3] B. Zhang, Y. Tian, and Z. Zhang. Network biology in medicine and beyond. Circulation: Cardiovascular Genetics, 7(4):536–547, 2014.

[4] C. Oates and S. Mukherjee. Network inference and biological dynamics. The annals of applied statistics, 6(3):1209, 2012.

[5] D. Koschutzki and F. Schreiber. Centrality analysis methods for biological networks and their application to gene regulatory networks. Gene regulation and systems biology, 2: GRSB–S702, 2008.

[6] E. Macau. A mathematical modeling approach from nonlinear dynamics to complex systems, volume 22. Springer, 2018.

[7] D. Koschutzki, H. Schwobbermeyer, and F. Schreiber. Ranking of network elements based on functional substructures. Journal of theoretical biology, 248(3):471–479, 2007.

[8] B. De Bivort, S. Huang, and Y. Bar-Yam. Empirical multiscale networks of cellular regulation. PloS computational biology, 3(10): e207, 2007.

[9] P. Langfelder and S. Horvath. “WGCNA: an R package for weighted correlation network analysis.” BMC bioinformatics 9.1 (2008): 1–13.

[10] W. Saelens, R. Cannoodt, and Y. Saeys. A comprehensive evaluation of module detection methods for gene expression data. Nature communications, 9(1):1–12, 2018

[11] S. Van Dam, U. Vosa, A. van der Graaf, L. Franke, and J. de Magalhaes. Gene coexpression analysis for functional classification and gene–disease predictions. Briefings in bioinformatics, 19(4):575–592, 2018.

[12] J. Liu, Z. Huole, and Q. Jianfeng. “Locally Adjust Networks Based on Connectivity and Semantic Similarities for Disease Module Detection.” Frontiers in genetics (2021): 1948.

[13] A. Werhli, M. Grzegorczyk, and D. Husmeier. Comparative evaluation of reverse engineering gene regulatory networks with relevance networks, graphical gaussian models and bayesian networks. Bioinformatics, 22(20):2523–2531, 2006.

[14] K. Murphy, S. Mian, et al. Modelling gene expression data using dynamic bayesian networks. Technical report, Citeseer, 1999.

[15] R. Steuer, J. Kurths, C. Daub, J. Weise, and J. Selbig. The mutual information: detecting and evaluating dependencies between variables. Bioinformatics, 18(suppl 2): S231–S240, 2002.

[16] S. Christley, N. Qing, and X. Xie. “Incorporating existing network information into gene network inference.” PloS one 4.8 (2009): e6799.

[17] Siahpirani, Alireza F., and Sushmita Roy. “A prior-based integrative framework for functional transcriptional regulatory network inference.” Nucleic acids research 45.4 (2017): e21–e21.

[18] Y. Li and J. Scott. “Gene network reconstruction by integration of prior biological knowledge.” G3: Genes, Genomes, Genetics 5.6 (2015): 1075–1079.

[19] X. Zhou and X. Cai. Inference of differential gene regulatory networks based on gene expression and genetic perturbation data. Bioinformatics, 36(1):197–204, 2020.

[20] J. Tu, L. Ou-Yang, Y. Zhu, H. Yan, H. Qin, and X. Zhang. Differential network analysis by simultaneously considering changes in gene interactions and gene expression. Bioinformatics, 2021

[21] D. Nitsch et al. Network analysis of differential expression for the identification of disease-causing genes. PloS one, 4(5): e5526, 2009.

[22] E. Macau. A mathematical modeling approach from nonlinear dynamics to complex systems, volume 22. Springer, 2018.

[23] D. Koschutzki, H. Schwobbermeyer, and F. Schreiber. Ranking of network elements based on functional substructures. Journal of theoretical biology, 248(3):471–479, 2007.

[24] F. Khawaja, et al. “Uncovering hidden community structure in multi-layer networks.” Applied Sciences 11.6 (2021): 2857.

[25] K. He, et al. “Revealing multiple layers of hidden community structure in networks.” arXiv preprint arXiv:1501.05700 (2015).

[26] L. Kogelman, et al. “Identification of co-expression gene networks, regulatory genes and pathways for obesity based on adipose tissue RNA Sequencing in a porcine model.” BMC medical genomics 7.1 (2014): 1–16.

[27] V. Huynh-Thu, et al. >“Inferring regulatory networks from expression data using tree-based methods.” PloS one 5.9 (2010): e12776.

[28] Huynh-Thu, Vân Anh, and Pierre Geurts. “dynGENIE3: dynamical GENIE3 for the inference of gene networks from time series expression data.” Scientific reports 8.1 (2018): 1–12.

[29] J. Vandesompele. “Accurate normalization of real-time quantitative RT-PCR data by geometric averaging of multiple internal control genes.” Genome biology 3.7 (2002): 1–12.

[30] V. Kumar and M. Sonajharia. “Feature selection: a literature review.” SmartCR 4.3 (2014): 211–229.

[31] Y. Saeys, I. Inaki, and L. Pedro. “A review of feature selection techniques in bioinformatics.” bioinformatics 23.19 (2007): 2507–2517.

[32] M. Love, S. Anders, and W. Huber. “Differential analysis of count data–the DESeq2 package.” Genome Biol 15.550 (2014): 10–1186.

[33] M. Zhang and Z. Zhou. “A k-nearest neighbor-based algorithm for multi-label classification.” 2005 IEEE international conference on granular computing. Vol. 2. IEEE, 2005.

[34] M. Newman. “Modularity and community structure in networks.” Proceedings of the national academy of sciences 103.23 (2006): 8577–8582.

[35] N. Dugué and P. Anthony. Directed Louvain: maximizing modularity in directed networks. Diss. Université d’Orléans, 2015.

[36] E. Leicht and M. Newman. “Community structure in directed networks.” Physical review letters 100.11 (2008): 118703.

[37] P. Hell and N. Jaroslav. “The core of a graph.” Discrete Mathematics 109.1-3 (1992): 117–126.

[38] L. Amaral, et al. “Classes of small-world networks.” Proceedings of the national academy of sciences 97.21 (2000): 11149–11152.

[39] M. Newman. “The structure and function of complex networks.” SIAM review 45.2 (2003): 167–256.

[40] M. Watson. “CoXpress: differential co-expression in gene expression data.” BMC bioinformatics 7.1 (2006): 1–12.

[41] S. Liu, et al. “OXPHOS deficiency activates global adaptation pathways to maintain mitochondrial membrane potential.” EMBO reports 22.4 (2021): e51606.

[42] M. Kuleshov, et al. “Enrichr: a comprehensive gene set enrichment analysis web server 2016 update.” Nucleic acids research 44.W1 (2016): W90–W97.

[43] A. Mordas and T. Kostas. “The MIA pathway: a key regulator of mitochondrial oxidative protein folding and biogenesis.” Accounts of chemical research 48.8 (2015): 2191–2199.

[44] F. Freimoser, et al. “Systematic screening of polyphosphate (poly P) levels in yeast mutant cells reveals strong interdependence with primary metabolism.” Genome biology 7.11 (2006): 1–9.

[45] A. Murari, et al. “IDH2-mediated regulation of the biogenesis of the oxidative phosphorylation system.” Science Advances 8.19 (2022): eabl8716.

[46] JC. Varela, et al. “The Saccharomyces cerevisiae HSP12 gene is activated by the high-osmolarity glycerol pathway and negatively regulated by protein kinase A.” Molecular and cellular biology 15.11 (1995): 6232–6245.

[47] P. Young, et al. “Activity-regulated cytoskeleton-associated protein (Arc/Arg3. 1) is transiently expressed after heat shock stress and suppresses heat shock factor 1.” Scientific reports 9.1 (2019): 1–13.

[48] Jaimes-Miranda, Fabiola, and Ricardo A. Chávez Montes. “The plant MBF1 protein family: a bridge between stress and transcription.” Journal of experimental botany 71.6 (2020): 1782–1791.

[49] M. Gerards, et al. “Intracellular vesicle trafficking plays an essential role in mitochondrial quality control.” Molecular biology of the cell 29.7 (2018): 809–819.

[50] H. Öhrvik, et al. “Ctr2 regulates biogenesis of a cleaved form of mammalian Ctr1 metal transporter lacking the copper-and cisplatin-binding ecto-domain.” Proceedings of the National Academy of Sciences 110.46 (2013): e4279–E4288.

[51] J. Ross, Jeremy., N. Zsuzsanna, and K. Robert. “The PHB1/2 phosphocomplex is required for mitochondrial homeostasis and survival of human T cells.” Journal of Biological Chemistry 283.8 (2008): 4699–4713.

[52] Yeo, Janet HC, et al. “A role for the mitochondrial protein Mrpl44 in maintaining OXPHOS capacity.” PloS one 10.7 (2015): e0134326.

[53] Boldogh, Istvan R., Hyeong-Cheol Yang, and Liza A. Pon. “Mitochondrial inheritance in budding yeast.” Traffic 2.6 (2001): 368–374.

[54] X. Xu, Xiuling, and F. Jeff. “A Role for the Transcriptional Coactivator PRC1 in Mitochondrial Biogenesis During Erythroid Development.” (2009): 3642–3642.

