## Supplementary material for "CoVar: A generalizable machine learning approach to identify the coordinated regulators driving variational gene expression": CoVar - Supplementary Method.docx

**Supplementary 4**

*1. Nearest neighbor network.*

Nearest neighbor (NN) is a machine learning classification technique in which a data point is assigned a class based on the class membership of majority of its most similar (or nearest) data point [1]. CoVar employs NN to identify the genes sharing the strongest interactions with the variational genes and constructs the nearest neighbor network constituting the variational genes, their neighbors, and their shared directed links. We present the following variants in NN construction.

*Coverage Approach.* For a variational gene, CoVar selects the top Z percentile nodes neighbor nodes each with the highest incoming and outgoing link weights in the perturbed network. Create a NN network of the variational genes and the nearest neighbors.

*Greedy Approach.* Unlike the coverage approach that assigns equal importance to all the variational genes, this approach selects the neighbors with the highest incoming and outgoing link weights across all the variational genes. It initializes the NN network with variational genes of the perturbed network, iteratively adds the nodes with the highest incoming or outgoing link weight to the existing NN network and stops when there are Z links in NN network.

It is noteworthy that unlike the GENIE perturbed and control networks, the NN network is not fully connected. It only possesses the highest link weights among the variational genes and their neighbors.

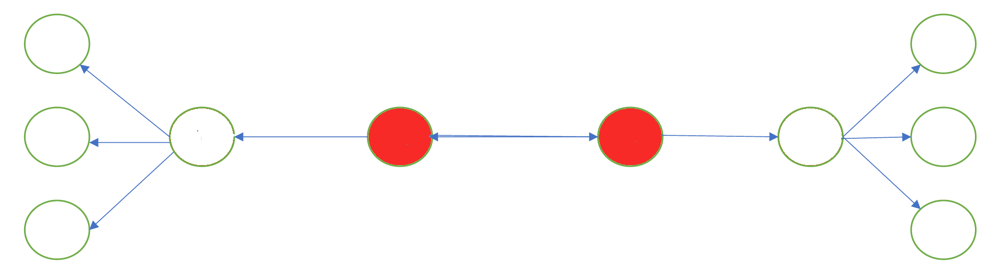

Figure 1. Identification of core nodes (marked red) from the nearest neighbor network

*2. Core network.*

The final step in CoVar is the identification of the core nodes, which are a subset of genes from the nearest neighbor (NN) network with two important properties, namely coordination and reachability. The core nodes (marked red) form a dense module with one another suggesting mutual influence or coordination. At the same time, the core nodes collectively have paths connecting most of the nodes in NN network, i.e., reachability.

There are two variants of core detection, namely, undirected and directed. Depending on the choice of approach, CoVar creates the undirected or directed core network from NN network. In the undirected approach, each core gene u has degree $deg(u) \geq k$, i.e., connections with $k$ other core nodes; similarly, in the directed approach, a core node $u$ has in-degree $\deg^{-} (u) \geq k$ and out-degree $\deg^{+} (u) \geq l$. We select the core network $G_{k,l}$ (with coordination and reachability scores DG and RG, respectively), such that there is no other core network $G_{k^{'},l^{'}}$ (where $(k, l)$ , $(k', l')$) such that $DG' > DG$ and $RG' > RG$. Figure 1 shows the core nodes 2 and 3 (marked red) in a 10-node NN network.

**References**

[1] M. Zhang and Z. Zhou. A k-nearest neighbor-based algorithm for multi-label classification. In 2005 IEEE international conference on granular computing, volume 2, pages 718–721. IEEE, 2005.
